## Supplementary material for "Global distribution of microbial carrageenan foraging pathways reveals widespread latent traits within the genetic “dark matter” of ruminant intestinal microbiomes": Source data 1: Alpaca, Group.pdf

### DIET SHEET - Alpaca

calgaryzoo

| | |
|---|---|
| **Common Name:** | Domestic Alpaca |
| **Scientific Name:** | *Lama pacos domestic* |
| **Animal Name (s):** | Latte (Fawn), Chai (white), Pekoe (brown) |
| **Accession Number:** | 108955,109046/7 |
| **Sex:** | F, F, F |
| **DOB:** | 15-Jul-10/6-Aug-12/11-Aug-12 |
| **Target BW Range (kg):** | 72, 95, 95 |
| **Target Calories (kcal):** | 1.5-2.5%BW, 70:30 F:C |
| **Calories Provided:** | 1.81 (2.8% BW), 80:20 F:C |
| **Avg. Intake (asfed)** | Intake not measured |

**Diet:** Alpaca
**Section:** South America

**Standard diet for :** 1
**Total same species in Enclosure:** 0.3

**STANDARD DIET FOR:** **1** **ANIMAL** **DATE:** **1-Sep-23**

| Day | | | | | | | Food Type | Amount | Notes |
|---|---|---|---|---|---|---|---|---|---|
| M | T | W | R | F | Sa | Su | Summer Herbivore Pellets* | 315 g | 1 2/3 cup, Flash with water, sprinkle with suppl. |
| M | T | W | R | F | Sa | Su | Alpaca Mineral Supplement | 30 g | balanced to 1.5ppm Se |
| M | T | W | R | F | Sa | Su | Ranch Mixed Hay** | 1 flake | 1.5kg/flake. Up to 4 flakes for the grou[illegible] Mixed hay, 10-20% alfalfa (10% CP) |

NOVEMEMBER 2023 SWITCH TO MAZURI ALPACA MAINTENANCE

Notes: Diet is per animal
*Pellet consumption tends to fluctuate, refer to whiteboard in cedar barn kitchen for current amounts.
Pellets are offered in the am as a shifting incentive. Any pellets used for alpaca BTS should come out of daily rations.
One Alpaca (Pekoe) has a history of choking episodes, therefore always ensure pellets are moistened.

**When transitioning to a new mixed hay, always do this slowly over 12 - 14 days.

***FEED AS INDICATED. DO NOT ALTER DIET. IF CHANGE IS REQUIRED PROVIDE DETAILS IN DIET CHANGE REQUEST.***
