## Supplementary material for "Global distribution of microbial carrageenan foraging pathways reveals widespread latent traits within the genetic “dark matter” of ruminant intestinal microbiomes": Source data 1: Appendix-7-Calgary-Zoo-Winter-Herbivore-Pellets-–-Formulae-code-M800710 (1).pdf

Wetaskiwin Coop Assoc. LTD.
Stored Formula Report
Plant : 108 - WETASKIWIN - FP Pricing
Plant: 108 - WETASKIWIN - FP

| Formula Code | Description | Species Code | Batch Weight | Date Stored | Ver |
|---|---|---|---|---|---|
| M800710 | Calgary Zoo WINTER Herbivore | 32 | 1000.00 | 17/02/2015 | 2 |

| Code | Ingredient Name | Amount |
|---|---|---|
| BPPG | Ground Beet Pulp | 401.23 |
| DAF | ALF-DEHY-17.5 | 247.50 |
| LINPRO | Linpro | 95.00 |
| WPP | MILLRUN | 92.50 |
| WDDG | Wheat Dist. Grain 50:50 | 50.00 |
| OMEGAFLX | Milled Flax-steel cut | 35.50 |
| OFLAXOIL | ORGANIC FLAX OIL | 30.00 |
| RPS | Canola Meal | 30.00 |
| SLT | SALT | 9.68 |
| LIM | LIMESTONE | 3.00 |
| RVE | VIT E-50% ADS | 1.48 |
| CAP | DICAL PHOS-21% | 1.00 |
| MGO | MAG OX-56% | 1.00 |
| 999 | Selenium 1000 mg/kg ( | 0.84 |
| 111 | TM SUL-Pak | 0.68 |
| 333 | B VIT PAK - P | 0.20 |
| 222 | ADE VIT PAK-30 Natur | 0.17 |
| 334466 | Sheep TM Micro Lt........ | 0.15 |
| D3BLEND | Vit D 10,000 IU/kg | 0.08 |

| No. | Nutrient Name | Units | Actual |
|---|---|---|---|
| 2 | Protein | % | 14.9507 |
| 3 | Fat | % | 7.8097 |
| 23 | linoleic acid | % | 2.9948 |
| 24 | linolenic acid | % | 0.6042 |
| 41 | TDN-ruminant | % | 72.4463 |
| 44 | NE maint | mcal/kg | 1.6784 |
| 50 | ME poultry | kcal/kg | 2,390.3750 |
| 56 | ME swine | kcal/kg | 2,610.8620 |
| 70 | calcium | % | 0.8315 |
| 71 | phosphorus | % | 0.3487 |
| 72 | av phosphorus | % | 0.1912 |
| 78 | magnesium | % | 0.3316 |

Wetaskiwin Coop Assoc. LTD.
Stored Formula Report
Plant : 108 - WETASKIWIN - FP Pricing
Plant: 108 - WETASKIWIN - FP

| No. | Nutrient Name | Units | Actual |
|---|---|---|---|
| 79 | potassium | % | 1.1510 |
| 81 | sodium | % | 0.5003 |
| 82 | sulfur | % | 0.1545 |
| 90 | cobalt | mg/kg | 0.7350 |
| 91 | copper | mg/kg | 17.0182 |
| 93 | iodine | mg/kg | 2.2875 |
| 94 | iron | mg/kg | 158.2434 |
| 95 | manganese | mg/kg | 135.8523 |
| 98 | zinc | mg/kg | 121.0324 |
| 110 | selenium added | mg/kg | 0.8400 |
| 116 | vit A | KIU/kg | 5.1000 |
| 117 | vit D3 | KIU/kg | 1.3100 |
| 118 | vit E | IU/kg | 754.6800 |
| 119 | vit K | mg/kg | 2.0000 |
| 120 | menadione | mg/kg | 2.0000 |
| 122 | biotin | mg/kg | 0.3374 |
| 123 | choline | mg/kg | 1,025.6120 |
| 124 | folic acid | mg/kg | 1.9742 |
| 125 | niacin | mg/kg | 53.7980 |
| 126 | pantothenic acid | mg/kg | 16.3615 |
| 127 | pyridoxine | mg/kg | 4.8580 |
| 128 | riboflavin | mg/kg | 9.4146 |
| 129 | thiamine | mg/kg | 3.8966 |
| 130 | vit B12 | mcg/kg | 16.0027 |
| 134 | vit A added | KIU/kg | 5.1000 |
| 135 | vit D3 added | KIU/kg | 1.3100 |
| 136 | vit E added | IU/kg | 754.5000 |
| 137 | vit K added | mg/kg | 2.0000 |
| 138 | menadione added | mg/kg | 2.0000 |
| 140 | biotin added | mg/kg | 0.2400 |
| 142 | folic acid added | mg/kg | 0.9000 |
| 143 | niacin added | mg/kg | 24.0000 |
| 144 | pantothenic acid ad | mg/kg | 7.0000 |
| 145 | pyridoxine added | mg/kg | 2.2000 |
| 146 | riboflavin added | mg/kg | 5.6000 |
| 147 | thiamine added | mg/kg | 1.3000 |
| 148 | vit B12 added | mcg/kg | 16.0000 |
