## Supplementary material for "Global distribution of microbial carrageenan foraging pathways reveals widespread latent traits within the genetic “dark matter” of ruminant intestinal microbiomes": Source data 1: Boar, European Wild, Adult, Female.pdf

### European Wild Boar, Female

| | |
|---|---|
| **Common Name:** | Wild Boar |
| **Scientific Name:** | *Sus scrofa scrofa* |
| **Animal Name (s):** | Fern, Poppy |
| **Accession Number:** | 109140/109141 |
| **Sex:** | F/F |
| **DOB:** | 3-Apr-13/15-Apr-13 |
| **Target BW Range (kg):** | 160 |
| **Target Calories:** | 2XME |
| **Calories Provided:** | |
| **Avg. Intake (asfed)** | 2.3, 1.4% BW |

**Diet:** Adult Female
**Section:** Asia

**Standard diet for :** 1
**Total same species in Enclosure:** 1.2

**STANDARD DIET FOR:** **1** **ANIMAL** **DATE:** **4-Aug-22**

| Day | | | | | | | Food Type | Amount | Notes |
|---|---|---|---|---|---|---|---|---|---|
| M | T | W | R | F | Sa | Su | Calgary Zoo Herbivore Cubes | 1500 g | |
| M | T | W | R | F | Sa | Su | Enrichment Fruit | 150 g | 40% apple, remove large pits |
| M | T | W | R | F | Sa | Su | Enrichment vegetables | 400 g | 50% Yam |
| M | T | W | R | F | Sa | Su | Yam | 250 g | |
| M | | W | | F | | | Greenie Dental Sticks - Large* | 1 stick | |

Preference list:
NO: Peppers of any color, NO carrots, NO celery
NO: turnip. NO cabbage, NO eggplant
NO: Orange, lemon, lime, or pinneapple

Notes: Diet is per individual. Monitor weights carefully, animals are prone to obesity and need to lose weight slowly.

***FEED AS INDICATED. DO NOT ALTER DIET. IF CHANGE IS REQUIRED PROVIDE DETAILS IN DIET CHANGE REQUEST.***

Recent changes:

*July-2022 begin trialing Greenie Dental Care 5" sticks for Large dog breeds to help with the chalky tartar build up vets noticed on their cheek teeth. Will reasses at next anesthetic event as these teeth are pretty deep in the mouth or in 6 months time (Jan 2023).
* Sept 26, 2023 - I wanted to note that Fern's teeth looked great during her immobilization today with regards to there being minimal gingivitis or tartar/calculus build up. Hopefully the greenies continue to help - ST

**Changes as of 12-July-2022**
Started offering dental sticks for dogs (4 inch, large dog breed). Barb did not want to try XL size.
2-3x per week, and will reasses at next anesthesia event because the teeth are back behind cheeks,

and hard to see without a knockdown
Barb and team will monitor to ensure they are chewed with their cheek teeth for at least 60 seconds or else may be poinless.

**Changes as of Spring 2022**

Due to HPAI strain, boars were kept inside, and received cracked corn to keep them occupied they all gained weight after this based on vet exam.
Also vets determined a lot of tartar build up from anesthesia event, trial dental sticks for dogs
