## Supplementary material for "Global distribution of microbial carrageenan foraging pathways reveals widespread latent traits within the genetic “dark matter” of ruminant intestinal microbiomes": Source data 1: Camel, Bactrian, Female.pdf

calgaryzoo

| | |
|---|---|
| Common Name: | Bactrian camel |
| Scientific Name: | *Camelus bactrianus* |
| Animal Name (s): | Zsa-Zsa |
| Accession Number: | 107222 |
| Sex: | F |
| DOB: | 23-Jun-05 |
| Target BW Range: | 750 (F) |
| Target Calories: | 1.5% BW |
| Calories Provided: | 1.12-1.3% BW |
| Avg. Intake (asfed) | |

**Diet:** Maintenance

**Section:** Asia

**Standard diet for :** 1

**Total same species in Enclosure:** 0.1

**STANDARD DIET FOR:** **1** **ANIMAL** **DATE:** **17-Sep-23**

| Day | | | | | | | Food Type | Amount | Notes |
|---|---|---|---|---|---|---|---|---|---|
| M | T | W | R | F | Sa | Su | Herbivore cubes (Zsa-Zsa) | 1500 g | 1 scoop |
| M | T | W | R | F | Sa | Su | Mixed hay (10-20% alfalfa, 10-13% CP) | 5-6 flakes | 1.5kg/flake |
| | T | | | | Sa | | Carrots | 125 g each | |
| | | | R | | | | Apples | 75 g each | |
| M | T | W | R | F | Sa | Su | Blue Salt Block | free choice | |
| M | T | W | R | F | Sa | Su | Browse | as available | low priority species |
| M | T | W | R | F | Sa | Su | Herbivore cubes | 200 g each | 1/2 AM 1/2 PM enrichment, meds |

Notes: Diet is per animal.
Low quality hay is sufficient, if too high in protein or alfalfa content, transition over 12-15 days, and may reduce amount offered. Assess BCS 2x per year.

***FEED AS INDICATED. DO NOT ALTER DIET. IF CHANGE IS REQUIRED PROVIDE DETAILS IN DIET CHANGE REQUEST.***
