## Supplementary material for "Global distribution of microbial carrageenan foraging pathways reveals widespread latent traits within the genetic “dark matter” of ruminant intestinal microbiomes": Source data 1: Caribou, Group.pdf

### DIET SHEET - Caribou

| | |
|---|---|
| **Common Name:** | American Woodland Caribou |
| **Scientific Name:** | *Rangifer tarandus caribou* |
| **Animal Name (s):** | Vanilla, Bean (Mica), Primrose, Avens |
| **Accession Number:** | 109193, 111023, 111327, X |
| **Sex:** | F, F, F, F |
| **DOB:** | 22-May-14, 29-Jun-20, 16-Jun-21, X |
| **Target BW Range (kg):** | 125 (F), 150 (M) |
| **Target Calories (kcal/d):** | 1.5-2.5% BW |
| **Calories Provided (kcal/d):** | 1.5-2.5% BW |
| **As fed, DM (kg/d), (%BW):** | F (1.9-3.1kg/d), M (2.25-3.75kg/d) |

**Diet:** Caribou
**Section:** Canadian Wilds

**Standard diet for :** 1
**Total same species in Enclosure:** 0.4

**STANDARD DIET FOR: 1 ANIMAL** **DATE: 1-Sep-23**

| Day | | | | | | | Food Type | Amount | Notes |
|---|---|---|---|---|---|---|---|---|---|
| M | T | W | R | F | Sa | Su | Winter Herbivore Pellets* | 1.5-2 rations per female | 1 ration = 1500g |
| M | T | W | R | F | Sa | Su | Winter Herbivore Pellets* | 2-3 rations per male | 1 ration = 1500g |
| M | T | W | R | F | Sa | Su | Alfalfa hay | free choice round bale or 2 flakes/d | |
| M | T | W | R | F | Sa | Su | Cobalt blue salt lick | free choice | 2kg size |
| M | T | W | R | F | Sa | Su | Lichen | as available | |
| M | T | W | R | F | Sa | Su | Approved Browse | free choice | Daily, 1st priority species<br>750g/d minimum 5% of forage |

LACTATING/NURSING ANIMALS

| Day | | | | | | | Food Type | Amount | Notes |
|---|---|---|---|---|---|---|---|---|---|
| M | T | W | R | F | Sa | Su | Calf Manna Pro | 200 g | Supplement to nursing animals |
| M | T | W | R | F | Sa | Su | Winter Herbivore Pellets | 1.5-2x current ration | |

Notes: Diet is per animal.

* Winter Herbivore Pellet fed year round, 1 ration = 1500g. When group fed, adjust up or down by one ration as needed. Only make 1 change per week, record in daily records.

When fed individually, increase or decrease by (10%) 150g as needed. Only make one change per week.

This species must be offered fresh or forzen browse daily

***FEED AS INDICATED. DO NOT ALTER DIET. IF CHANGE IS REQUIRED PROVIDE DETAILS IN DIET CHANGE REQUEST.***
