## Supplementary material for "Global distribution of microbial carrageenan foraging pathways reveals widespread latent traits within the genetic “dark matter” of ruminant intestinal microbiomes": Source data 1: Giraffe, Adult, Female.pdf

### Giraffe, Female

| | |
|---|---|
| Common Name: | Giraffe |
| Scientific Name: | *Giraffa camelopardalis* |
| Animal Name (s): | Moshi, Emara |
| Accession Number: | 109513, 109615 |
| Sex: | F |
| DOB: | Oct. 2015, May 2011 |
| Target BW Range: | 850 |
| Target Calories: | 2% BW |
| Calories Provided: | 2 %BW |
| Avg. Intake (asfed, kg) | 17 |

| | |
|---|---|
| Diet: | Maintenance |
| Section: | Savannah |
| Standard diet for : | 1 |
| Total same species in Enclosure: | 1.2 |

**STANDARD DIET FOR: 1 ANIMAL DATE: 15-Oct-22**

| Day | | | | | | | Food Type | Amount | Notes |
|---|---|---|---|---|---|---|---|---|---|
| M | T | W | R | F | S | Su | Winter Herbivore Pellets | 3200 g | Moshi - 2900g (23-Jan-2021) |
| M | T | W | R | F | S | Su | Alfalfa Hay | 8000 g | |
| M | T | W | R | F | S | Su | Vegetable Variety | 200 g | |
| M | T | W | R | F | S | Su | Fruit Variety | 50 g | |
| M | T | W | R | F | S | Su | Cobalt Blue Salt Block | free choice | |
| M | | W | | F | | | Primate Biscuits | 1000 g | Training, hoof trims, etc |
| | | | | | | Su | Carrots, Lettuce | 1000 g | Zoo School, BTS, 1-4x per month |
| M | T | W | R | F | S | Su | Browse (Fresh, Frozen, or Silage) | 5000 g | 1st priority species<br>5% of forage, minimum requirement |

Notes: Diet is per individual.
Winter Herbivore Pellets all year round.
Extra bananas reserved for medicating, not included in diet
Recommended 16-20% CP, 70:30 F:C

***FEED AS INDICATED. DO NOT ALTER DIET. IF CHANGE IS REQUIRED PROVIDE DETAILS IN DIET CHANGE REQUEST.***
