## Supplementary material for "Global distribution of microbial carrageenan foraging pathways reveals widespread latent traits within the genetic “dark matter” of ruminant intestinal microbiomes": Source data 1: Giraffe, Adult, Male.pdf

### Giraffe, Male

| | |
|---|---|
| **Common Name:** | Giraffe |
| **Scientific Name:** | *Giraffa camelopardalis* |
| **Animal Name (s):** | Nabo |
| **Accession Number:** | 109025 |
| **Sex:** | M |
| **DOB:** | 1-Jan-10 |
| **Target BW Range (kg):** | 1150 |
| **Target Calories:** | 2% BW |
| **Calories Provided:** | 2 %BW |
| **Avg. Intake (asfed, kg)** | 22.7 |

| | |
|---|---|
| **Diet:** | Maintenance |
| **Section:** | Savannah |
| **Standard diet for :** | 1 |
| **Total same species in Enclosure:** | 1.2 |

**STANDARD DIET FOR: 1 ANIMAL** **DATE: 6-Aug-22**

| Day | | | | | | | Food Type | Amount | Notes |
|---|---|---|---|---|---|---|---|---|---|
| M | T | W | R | F | S | Su | Winter Herbivore Pellets | 3400 g | |
| M | T | W | R | F | S | Su | Alfalfa Hay | 12000 g | |
| M | T | W | R | F | S | Su | Vegetable Variety | 200 g | |
| M | T | W | R | F | S | Su | Fruit Variety | 50 g | |
| M | T | W | R | F | S | Su | Cobalt Blue Salt Block | free choice | |
| M | | W | | F | | | Primate Biscuits | 1000 g | Training, hoof trims, etc |
| | | | | | | Su | Carrots, Lettuce | 1000 g | Zoo School, BTS, 1-4x per month |
| M | T | W | R | F | S | Su | Browse (Fresh, Frozen, or Silage) | 6500 g | 1st priority species<br>5% of forage, minimum requirement |

Notes: Diet is per individual.
Winter Herbivore Pellets all year round.
Recommended 16-20% CP, 70:30 F:C

***FEED AS INDICATED. DO NOT ALTER DIET. IF CHANGE IS REQUIRED PROVIDE DETAILS IN DIET CHANGE REQUEST.***
