## Supplementary material for "Global distribution of microbial carrageenan foraging pathways reveals widespread latent traits within the genetic “dark matter” of ruminant intestinal microbiomes": Source data 1: Goat, Rocky Mountain, Group.pdf

Wilder Institute Calgary Zoo

### Rocky Mountain Goat

| | |
|---|---|
| **Common Name:** | Rocky Mountain Goat |
| **Scientific Name:** | *Oreamnos americanus* |
| **Animal Name (s):** | Shannon, Amanda, Yukon, Peyto, Pika, Suncup, Hara |
| **Accession Number:** | 107724, 5, 109032, 110805, 6, 111113, 111486 |
| **Sex:** | F/F/M/F/F/F/F |
| **DOB:** | 23-May-07 (2), 29-May-12, 18-May-20, (2), 10-May-21, 10-May-22 |
| **Target BW Range (kg):** | 75-85 |
| **Target Calories:** | 2-3.5%BW, 60:40 F:C |
| **Calories Provided:** | |
| **Avg. Intake (asfed)** | 2.0-3.5% BW |

**Diet:** Maintenance

**Section:** Canadian Wilds

**Standard diet for :** 1

**Total same species in Enclosure:** 1.6

**STANDARD DIET FOR: 1 ANIMAL** **DATE: 3-Aug-22**

| Day | | | | | | | Food Type | Amount | Notes |
|---|---|---|---|---|---|---|---|---|---|
| M | T | W | R | F | S | Su | Calgary Zoo Herbivore Pellets* | 1100 g | 7700 g for group maintenance** |
| M | T | W | R | F | S | Su | Mixed Hay (25% alfalfa) | 2/3 flake | Grassy, soft, 10-20% alfalfa, 16-18%CP |
| M | T | W | R | F | S | Su | Cobalt Blue Salt Block | ad libitum | |
| | | | | | S | | Romaine | 0.25 hd | 1.5hd for group |
| M | T | W | R | F | S | Su | Browse | as available | 2nd priority species<br>2.5% of forage, 300g/d |
| | | | | | | | Edible Flowers | as available | seasonal, by donation |

Notes:
Diet is per animal.
*Summer herbivore pellet April to Oct. Transition over 12 days 25:75 (4 days), 50:50 (4 days), 75:25 (4 days), 100%
*Winter herbivore pellet Nov to Mar. Transition over 12 days 25:75 (4 days), 50:50 (4 days), 75:25 (4 days), 100%
Consult animal nutrition supervisor for current mixed hay components and recommended amounts.
Recommended, 25% Alfalfa, 16-18%CP, 90% DMB
Pasture available in summer, hay may be reduced based on appetite and body condition
**Adjust amount of herbivore pellets based on number of animals and stage of gestation (2x in 3rd trimester) or lactation (3x). Offer Foal Lac to goat kids.
Difficult to enrich with food, as they are very picky. Scent enrichment is recommended.

***FEED AS INDICATED. DO NOT ALTER DIET. IF CHANGE IS REQUIRED PROVIDE DETAILS IN DIET CHANGE REQUEST.***
