## Supplementary material for "Global distribution of microbial carrageenan foraging pathways reveals widespread latent traits within the genetic “dark matter” of ruminant intestinal microbiomes": Source data 1: Hippo.pdf

### Hippo, Adult

| | |
|---|---|
| Common Name: | Hippopotamus |
| Scientific Name: | *Hippopotamus amphibius* |
| Animal Name (s): | Lobi, Sparky |
| Accession Number: | 109024, 102964 |
| Sex: | M, F |
| DOB: | 31-Oct-06, 30-May-87 |
| Target BW Range (kg): | 1900, 1400 (est) |
| Est. Pred. Equ. (kcal/d): | 40290, 32042 |
| Calories Provided (kcal/d): | 21577 |
| Intake asfed, kg, %BW: | 14.1, 0.85 |

**Diet:** Maintenance
**Section:** Savannah

**Standard diet for :** 1
**Total same species in Enclosure:** 1.1

**STANDARD DIET FOR: 1 ANIMAL** **DATE: 9-Feb-21**

| Day | | | | | | | Food Type | Amount | | Notes |
|---|---|---|---|---|---|---|---|---|---|---|
| M | T | W | R | F | Sa | Su | Herbivore Cubes | 3000 g | each | |
| M | T | W | R | F | Sa | Su | Vitamin E (50%) | 2 tsp. | each | sprinkle on top of cubes |
| M | T | W | R | F | Sa | Su | Biotin (2%) | 1.5 tsp. | each | sprinkle on top of cubes |
| M | T | W | R | F | Sa | Su | Timothy hay | 4 flakes | each | ~ 6kg/day each |
| M | T | W | R | F | Sa | Su | Mixed Hay Alfalfa-grass hay | 1.5 2" flake | each | ~ 3kg/day each |
| M | T | W | R | F | Sa | Su | Romaine | 1600 g | 2 hd each | fed in pool at any time |
| M | T | W | R | F | Sa | Su | Fruit | 300 g | 1000g for 1.1, training/enrichment | |
| M | T | W | R | F | Sa | Su | Vegetables | 200 g | | |

Notes: Calories calculated using Kleiber 2XBMR (LJ)
4-May-2022 - Lobi's diet reduced to Adult ration (3500g to 3000g herb cubes over 4 weeks)

***FEED AS INDICATED. DO NOT ALTER DIET. IF CHANGE IS REQUIRED PROVIDE DETAILS IN DIET CHANGE REQUEST.***
