## Supplementary material for "Global distribution of microbial carrageenan foraging pathways reveals widespread latent traits within the genetic “dark matter” of ruminant intestinal microbiomes": Source data 1: Markhor, Group.pdf

| | |
|---|---|
| Common Name: | Markhor |
| Scientific Name: | *Capra falconeri* |
| Animal Name (s): | Sproing, Cliff |
| Accession Number: | 107234, 111490 |
| Sex: | F, M |
| DOB: | 12-May-05, 17-Aug-21 |
| Target BW Range: | 48 (F), growth (M) |
| Target Calories: | 1.5-2.5% BW |
| Calories Provided: | |
| Avg. Intake (asfed) | 1.8-4% BW |

**Diet:** Maintenance

**Section:** ASIA

**Standard diet for :** 1

**Total same species in Enclosure:** 1.1

**STANDARD DIET FOR:** 1 **ANIMAL** **DATE:** **17-Sep-23**

| Day | | | | | | | Food Type | Amount | g | Group (1.1) Amount |
|---|---|---|---|---|---|---|---|---|---|---|
| M | T | W | R | F | Sa | Su | Winter Herbivore Pellet | 1.5-3 cups | 265-530 | 3-6 cups |
| M | T | W | R | F | Sa | Su | Wild Herbivore Plus | 0.75-1.5 cups | 125-250 | 1.5-3 cups |
| | | | | | | | 2:1 ratio winter:WH+ | | | |
| M | T | W | R | F | Sa | Su | Mixed Hay (>50% alfalfa) | 1/4-1/2 flakes | 375-750 | 1.5-2 flakes |
| | T | | | | Sa | | Romaine | 1/4 hd | 200 | 1/2 hd |
| | | W | | F | | Su | Apple or Carrot | 50 g | | 100 g |
| M | T | W | R | F | Sa | Su | Cobalt Blue Salt Block | free choice | | |
| M | T | W | R | F | Sa | Su | Browse | as available | | 2nd priority browse species min 225g/day, 2.5% of forage |

Notes: Diet is per animal. Group amount shown for 1.1.0

Hay and Pellet intake changes seasonally and is recorded daily on section, adjusted as needed.

Feed high quality hay (>50% alfalfa, CP>14%) over the winter (Nov to June), may reduce quality July to Oct (<50% alfalfa, CP 11-13%).

Remain on Winter Herb Pellet and WH+ year round

Produce is used for training and enrichment purposes, or medicating.

Breeding Season Jan (rut) - April, may reduce appetite and BCS.

Pregnant animals in 3rd trimester or lactating animals may receive upto 30% increase in pellet and hay components.

BCS assessed quartely (May/Jun, Aug/Sept, Nov/Dec, and Feb/Mar)

Offer boost at 10% of diet for thin/geriatic animals (75-150g/d).

***FEED AS INDICATED. DO NOT ALTER DIET. IF CHANGE IS REQUIRED PROVIDE DETAILS IN DIET CHANGE REQUEST.***
