## Supplementary material for "Global distribution of microbial carrageenan foraging pathways reveals widespread latent traits within the genetic “dark matter” of ruminant intestinal microbiomes": Source data 1: Mazuri Mini Leaf-Eater.pdf

Mazuri®
EXOTIC ANIMAL NUTRITION

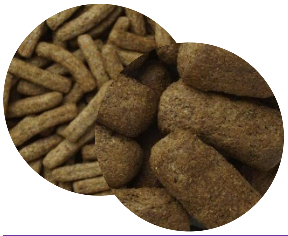

### Leaf-Eater Primate Diets

Mazuri® Leaf-Eater Primate Diets are complete lifecycle diets specially formulated to meet the needs of leaf-eating primates which are thought to require a high-fiber diet such as lemurs, langurs, and howlers. This diet can be fed to other species of primates such as gorillas and orangutans, when a high-fiber diet is desired.

#### Features and Benefits

- **Designed to be fed with supplementation** – Allows for natural feeding behaviors by the addition of species-appropriate food items.
- **Meets NRC recommendations for all nutrients** – Except protein, sodium, and chloride when fed at 50% of the diet.
- **High fiber** – Contains multiple sources of soluble and insoluble fiber.
- **No added sucrose or fructose** – Helps maintain dental health and may be appropriate for sugar-sensitive primates.
- **No added wheat product** – May be appropriate for animals with wheat sensitivity.
- **Enriched with Vitamins E and vitamin C** – Supports animal wellness.
- **Contains flaxseed** – Source of Omega-3 fatty acids.

#### Product Form & Packaging

Extruded Particle | 25 lb. (11.33 kg) net weight paper sack

- **Catalog #0001472** | 1” x 2” Biscuit
- **Catalog #0001448** | 1/4” x 1” Mini biscuit

#### Guaranteed Analysis

| | | | |
|---|---|---|---|
| Crude protein not less than | 23.00% | Moisture not more than | 12.00% |
| Crude fat not less than | 5.00% | Ash not more than | 9.00% |
| Crude fiber not more than | 14.00% | | |

#### Ingredients

Dehulled Soybean Meal, Ground Soybean Hulls, Ground Corn, Corn Gluten Meal, Ground Oats, Dried Plain Beet Pulp, Dried Apple Pomace, Soybean Oil, Dehydrated Alfalfa Meal, Dicalcium Phosphate, Calcium Carbonate, Ground Flaxseed, Brewers Dried Yeast, Salt, L-Ascorbyl-2-Polyphosphate (Vitamin C), DL-Methionine, Pyridoxine Hydrochloride, Choline Chloride, Folic Acid, Vitamin A Acetate, Cholecalciferol (Vitamin D3), d-Alpha Tocopheryl Acetate (Vitamin E), Calcium Pantothenate, Ferrous Sulfate, Menadione Sodium Bisulfite Complex (Vitamin K), Preserved with Mixed Tocopherols, Manganous Oxide, Zinc Oxide, Rosemary Extract, Ferrous Carbonate, Nicotinic Acid, Citric Acid (a Preservative), Thiamine Mononitrate, Vitamin B12 Supplement, Riboflavin Supplement, Copper Sulfate, Zinc Sulfate, Calcium Iodate, Cobalt Carbonate, Sodium Selenite, Biotin.

#### Feeding Directions

- Mazuri® Primate Diets are designed to be an essential part of a total primate feeding system and may be used in conjunction with all other Mazuri® primate products.
- Primates generally consume 2% to 4% of their body weight in food each day on a dry matter basis (i.e., a 50 lb. animal will eat 1 to 2 kg of food per day, on a dry matter basis).
  - Mazuri® primate diets can be supplemented with fresh vegetables and fruit if this is desired to provide variety in the diet, as long as the dry matter of these items does not exceed 50% of the dry matter consumption. Using typical produce and non-toxic browse, a feeding program on an “as fed” basis might consist of 60% produce/browse and 40% Mazuri® primate products (i.e., a diet may be 6 lb. of produce/browse plus 4 lb. of one or a mix of Mazuri® primate products).
- The amount of diet the animal requires will vary according to its age, size, life stage, health status, and activity of the animal as well as the environmental temperature.
  - Adjust daily feed intake based on the health status, body condition, and nutrient level desired to meet the needs of your primates.
  - Diet may be soaked to soften the biscuits for neonates or animals that have difficulty chewing.
- Primates require ascorbic acid (vitamin C) in their daily diet. Mazuri® Primate Diets contain a stabilized form of vitamin C that is stable under appropriate storage conditions for at least 1 year from the date of manufacturing printed on the bag.
- Always provide plenty of fresh, clean water. Thoroughly wash feed and water bowls on a regular basis. It is always good practice to wash hands thoroughly after feeding and/or handling animals.
- This diet is not for human consumption.

#### Storage Conditions

For best results, reseal the bag between uses or store contents of open paper sack in container with sealing lid. Store in a cool (75°F/24°C or colder), dry (approximately 50% RH) location free from rodents and insects. Do not offer moldy or insect-infested feed to animals as it may result in illness, performance loss or death. Freezing will not harm the diet and may extend freshness. Use within 1 year of bag manufacturing or "Best if Used By" date.

Mazuri® is a registered trademark of PMI Nutrition International, LLC.
