## Supplementary material for "Global distribution of microbial carrageenan foraging pathways reveals widespread latent traits within the genetic “dark matter” of ruminant intestinal microbiomes": Source data 1: Moose.pdf

calgaryzoo

| | |
|---|---|
| **Common Name:** | Moose |
| **Scientific Name:** | *Alces americanus* |
| **Animal Name (s):** | Maple, Aspen |
| **Accession Number:** | 109978, 110242 |
| **Sex:** | F/F |
| **DOB:** | 26-May-18, 18-May-07 |
| **Target BW Range (kg):** | 250-300 |
| **Target Calories:** | 3-4% BW, 15-20%CP |
| **Calories Provided:** | 3.4% BW, 60:40 F:C |
| **Avg. Intake (asfed)** | seasonal |

| | |
|---|---|
| **Diet:** | Maintenance |
| **Section:** | Canadian Wilds |
| **Standard diet for :** | 1 |
| **Total same species in Enclosure:** | 0.2 |

**STANDARD DIET FOR:** **1** **ANIMAL** **DATE:** **1-Jul-22**

| Day | | | | | | | Food Type | Amount | Notes |
|---|---|---|---|---|---|---|---|---|---|
| M | T | W | R | F | Sa | Su | Mazuri Moose Maintenance | 3600 g | between 2500 and 4500g* |
| M | T | W | R | F | Sa | Su | alfalfa hay, prime quality | 250 g | seasonal intake |
| M | T | W | R | F | Sa | Su | Cobalt Blue Salt Block | ad libitum | |
| M | T | W | R | F | Sa | Su | Browse | as available | 1st priority species<br>2.5kg/day min, 5% of forage |

Notes: *Diet intake changes seasonally. Monitor intake and fecal scores daily.
Offer Moose Breeder to breeding animals or 50:50 (breeder: maintenance) to growing moose upto the age of 4, double intake during 3rd trimester and lactation.

***FEED AS INDICATED. DO NOT ALTER DIET. IF CHANGE IS REQUIRED PROVIDE DETAILS IN DIET CHANGE REQUEST.***
