## Supplementary material for "Global distribution of microbial carrageenan foraging pathways reveals widespread latent traits within the genetic “dark matter” of ruminant intestinal microbiomes": Source data 1: Musk Deer.pdf

| | |
|---|---|
| **Common Name:** | Siberian Musk Deer |
| **Scientific Name:** | *M. moschiferus* |
| **Animal Name (s):** | Ozzy |
| **Accession Number:** | 109476 |
| **Sex:** | M |
| **DOB:** | 12-Jun-13 |
| **Target BW Range (kg):** | 11-12 |
| **Target Calories (kcal/d):** | 891-1366 |
| **Calories Provided:** | 1047 |
| **Avg. Intake (asfed, kg)** | 0.56 (+ forage) |

**Diet:** Maintenance
**Section:** Asia

**Standard diet for :** 1
**Total same species in Enclosure:** 1.0

**STANDARD DIET FOR:** **1** **ANIMAL** **DATE:** **10-Oct-23**

| Day | | | | | | | Food Type | Amount | Notes |
|---|---|---|---|---|---|---|---|---|---|
| M | T | W | R | F | S | Su | Mazuri Mini Leaf-Eater | 137 g | 1, 1/4 cup |
| M | T | W | R | F | S | Su | Winter Herbivore Pellets | 132 g | 3/4 cup |
| M | T | W | R | F | S | Su | Yam | 60 g | chopped 2mmx2mmx2mm |
| M | T | W | R | F | S | Su | Alfalfa Hay | 1/6 flake | 2 large handfuls (250g), mostly leaf |
| M | T | W | R | F | S | Su | Cobalt Blue Salt block | 1 block | available all times |
| M | T | W | R | F | S | Su | Browse | 250 g | 2nd priority species<br>35g/d, 2.5% forage |

Notes: Diet is per animal per day.
Diet accounts for seasonal variation in intake and pests.

***FEED AS INDICATED. DO NOT ALTER DIET. IF CHANGE IS REQUIRED PROVIDE DETAILS IN DIET CHANGE REQUEST.***
