## Supplementary material for "Global distribution of microbial carrageenan foraging pathways reveals widespread latent traits within the genetic “dark matter” of ruminant intestinal microbiomes": Source data 1: Sheep, Bighorn, Group.pdf

| | |
|---|---|
| **Common Name:** | Bighorn Sheep |
| **Scientific Name:** | *Ovis canadensis* |
| **Animal Name (s):** | Callie, Bill Williams, Cheyenne, June, Cash |
| **Accession Number:** | 107661,109708/9,107962,110144 |
| **Sex:** | F, M, F, F, F |
| **DOB:** | 15-May-06, 3-May-17, 3-May-17, 4-Jun-18, 18-Jul-18 |
| **Target BW Range (kg):** | 35-90 |
| **Target Calories:** | 2-3.5%BW, 60:40 F:C |
| **Calories Provided:** | |
| **Avg. Intake (asfed)** | |

| | |
|---|---|
| **Diet:** | Maintenance |
| **Section:** | CW |
| **Standard diet for :** | 1 |
| **Total same species in Enclosure:** | 1.4 |

**STANDARD DIET FOR: 1 ANIMAL** **DATE: 3-Aug-22**

| Day | | | | | | | Food Type | Amount | Notes |
|---|---|---|---|---|---|---|---|---|---|
| M | T | W | R | F | S | Su | Calgary Zoo Herbivore Pellets* | 750 g | 3750g/group |
| M | T | W | R | F | S | Su | Mixed Hay (25% alfalfa) | 3/5 flake | 3 flakes/group, 4.5kg/day |
| M | T | W | R | F | S | Su | Cobalt Blue Salt Block | ad libitum | |
| | | | | | S | | Romaine | 0.25 hd | 1.5hd for group |
| M | T | W | R | F | S | Su | Browse | as available | 2nd priority species<br>250-500g/day, 2.5% of forage |
| | | | | | | | Edible Flowers | as available | seasonal, by donation |

Notes: Diet is per animal. This is a Copper sensitive species, highly susceptible to Cu toxicity.

*Summer herbivore pellet April to Oct. Transition over 12 days 25:75 (4 days), 50:50 (4 days), 75:25 (4 days), 100%

*Winter herbivore pellet Nov to Mar. Transition over 12 days 25:75 (4 days), 50:50 (4 days), 75:25 (4 days), 100%

Consult animal nutrition supervisor for current mixed hay components and recommended amounts.

Recommended, 25% Alfalfa, 15-18%CP, 90% DMB

Pasture available in summer., hay may be reduced based on appetite and body condition

Adjust amount of herbivore pellets based on number of animals and stage of gestation or lactation (currenlty not breeding, all animals are related)

Difficult to enrich with food, as they are very picky. Scent enrichment is recommended.

***FEED AS INDICATED. DO NOT ALTER DIET. IF CHANGE IS REQUIRED PROVIDE DETAILS IN DIET CHANGE REQUEST.***
