## Supplemental text for "Global distribution of microbial carrageenan foraging pathways reveals widespread latent traits within the genetic “dark matter” of ruminant intestinal microbiomes"

### Supplemental information

#### Supplemental note 1

#### Spatial digestion of *M. japonica* in bovine GIT

*Bacteroidota*, primarily *Prevotella*, are widely recognized as one of the most abundant fibre degrading bacterial phyla in the rumen[1, 2]. Often overlooked, however, is the prowess of *Bacteroides* within the hindgut of ruminants that forage on undigested feed residues that bypass the rumen[1], such as cellulose and hemicelluloses[3, 4]. This can be attributed to increased bulk within the digestive tract[39], which could be impacted by seaweed polysacchrides as observed in the digesta of mice[5]. Metagenome-guided metaproteomics analysis presented here suggests that *Bacteroides*-affiliated MAGs are one of the primary mechanisms for microbial degradation and utilization of *M. japonica* carrageenans in the lower GIT of cattle **(Fig. 1; Extended Data Fig. 2C)**. There was a minor increase in ruminal *Bacteroidaceae* under *M. japonica* supplementation, which paled in comparison to the increase in fecal samples. This is likely due to *Bacteroides* spp. being more abundant within the lower gastrointestinal tract[1] (**Extended Data Fig. 1A**). These findings suggest the disparity between research on the rumen microbiome compared to the lower GIT microbiome, may hamper the discovery of seaweed degrading pathways within ruminants. Further, the impact carrageenans, and potentially other seaweed polysaccharides, has on the lower GIT microbiome may provide targeted prebiotic benefits or function as delivery systems to the hindgut of ruminants.

#### Supplemental results

#### Confirmation of *M. japonica* well wall polysaccharides

Sulfated galactans were extracted from *M. japonica* and studied by linkage analysis via methylation-GC-MS analysis of PMAAs prepared from the native and desulfated products[6, 7]. Methylation-GC of the sulfated galactan without desulfation showed a high level of 3,4-Gal*p* at 38.1%, followed by 4-AnGal*p* (16.5 %), 2,4-AnGal*p* (14.4 %), 3-Gal*p* (12.1 %), 2,3-Gal*p* (7.1 %), 2,4,6-Gal*p* (3.1 %), 4,6-Gal*p* (2.5 %) (**Supplemental Fig. 1A**), and various other minor Gal*p* linkages (**Supplementary Table 17**). Compared to the sample without desulfation, the sample subjected to 6 h of solvolytic desulfation[8] was dominated by 3-Gal*p* (55.8 %), followed by 4-AnGal*p* (15.5 %), 4-Gal*p* (15.3 %), 3,4-Gal*p* (6.2 %), 3,6-Gal*p* (2.5 %), and other minor Gal*p* linkages (**Supplementary Table 17**). This was not appreciably different than desulfation at 16 h (**Supplemental Fig. 1B; Supplementary Table 17**). Glycosidic linkages of galactoses were confirmed based on the EI-MS ion fragmentation patterns of their PMAAs (**Supplemental Fig. 1C**), with reference to published patterns in the literature[9]. The results suggested that the majority of 3,4-Gal*p* was converted to 3-Gal*p* due to desulfation at the *O*-4 position (**Supplemental Fig. 1D**). A considerable amount of 4-Gal*p* was also detected in the desulfated galactan, indicating that the 4-linked galactose residues were sulfated as well. There was no 2,4-AnGal*p* was detected in the desulfated samples, indicating the complete conversion of 2,4-AnGal*p* to 4-AnGal*p* by desulfation at the *O*-2 position. The total level of AnGal in the desulfated sample was less than half of that in the untreated sample, indicating partial degradation of AnGal caused by solvolytic desulfation[8]. This was further supported by the lower level of anhydro sugar in the 16 h treatment compared to the 6 h one (**Supplementary Table 17**).

**_Bx_Car CarPUL %GC content is distinct _Bx_Car genomes**

Mean GC content percentage between the *Bx*Car$_{BOV}$ genomes and CarPULs was compared to that of control *Bx*XB1A$_{HOM}$, xylan PULs (terrestrial plant polysaccharide) from *Bx*Car5$_{BOV}$ and 17, and marine/sediment microorganisms: *L. citreus* and *S fermentans, L. marinus, Saccharicrinis aurantiacus, Flavobacterium sp. 7A,* and *Wenyingzhuangia aestuarii*. These were selected based on the high homology observed in CarPUL CAZyme BLAST results. The mean GC% from *Bx*CAR$_{BOV}$ CarPUL genes was more similar to those of marine pelagic and sediment organisms genes than to the remainder of the BxCAR$_{BOV}$ genome (**Supplemental Fig. 2**). Through Games Howell's multiple comparisons tests, the mean GC% of *Bx*Car5$_{BOV}$ CarPUL-1 (39.3 ± 4.8) was more closely aligned with the *S. fermentans* genome (37.7 ± 3.8$^{ns}$) than to itself (42.5 ± 5.3$^{****}$), a terrestrial xylan PUL[10] within the *Bx*Car5$_{BOV}$ genome (44.4 ± 5.0$^{**}$), and the reference *Bx*XB1A$_{HOM}$ genome (42.6 ± 5.0$^{****}$); this pattern is repeated between the *Bx*Car17$_{BOV}$ CarPUL and similar datasets. *Bx*Car5$_{BOV}$ CarPUL-2 (35.7 ± 3.6) showed the highest similarity towards marine pelagic and sediment organisms, with GC% values showing no significant difference to ORFs within *S. aurantiacus* (35.4 ± 3.4$^{ns}$)*, Flavobacterium* sp. 7A (34.3 ± 3.6$^{ns}$)*,* and *L. marinus* (35.6 ± 3.1$^{ns}$). All GC values and statistics are in **Source Data 2**.

### Supplemental figures

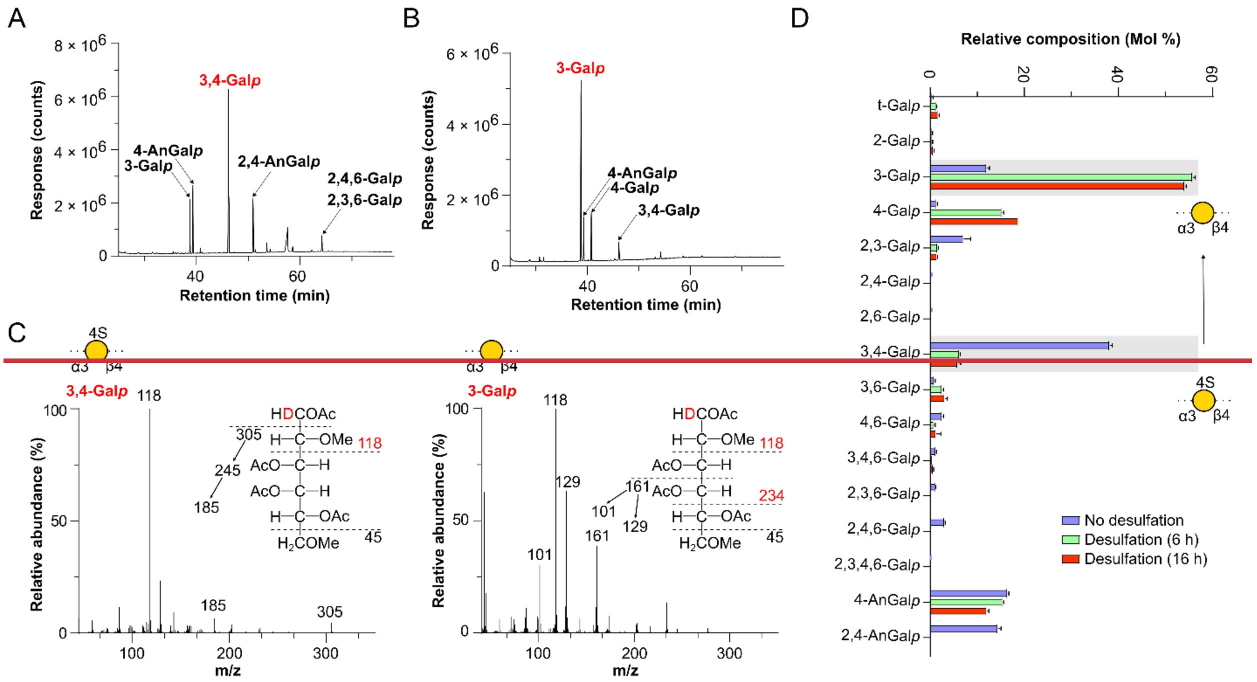


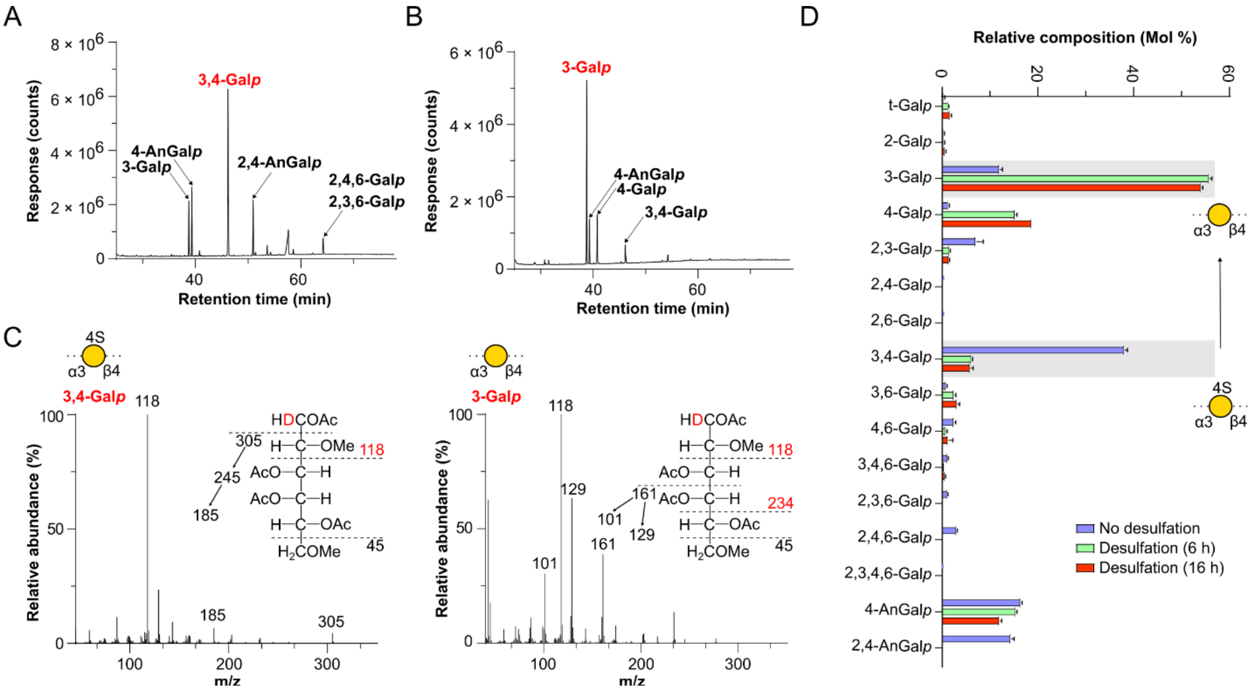


**Supplemental figure 1: Glycosidic linkage analysis (methylation-GC-MS analysis) of *M. japonica* galactan and its desulfation product.** GC-TIC chromatograms of PMAAs from destarched, water-soluble *M. japonica* extract (*Mj*Ex) **A)** before solvolytic desulfation and **B)** after 16 h of solvolytic desulfation. **C)** EI-MS spectra and ion fragmentation patterns of PMAAs from the largest peaks within GC-TIC chromatograms. Top: no desulfation (3,4-Galp). Bottom: 16 h desulfation (3-Galp). **D)** Comparison of relative composition (Mol%) of observed PMAAs from samples with no desulfation and those subjected to 6 h and16 h of desulfation.

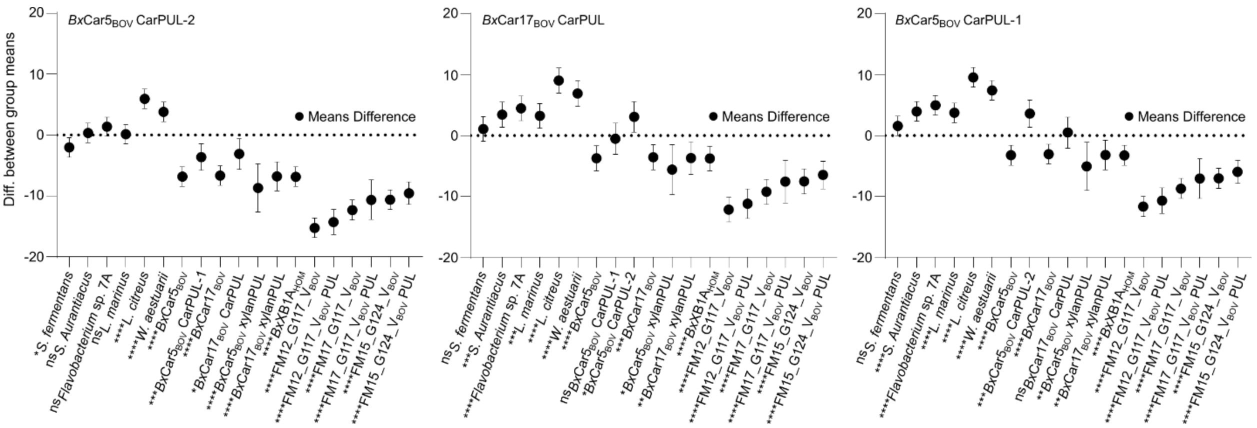


**Supplemental figure 2: Mean % GC content between *Bx*Car CarPULs and *Bx*Car genomes is significantly different**. Mean %GC content of CarPULs was compared to that of their genome (whole genome and terrestrial xylan PULs[10] within their genome), marine pelagic and sediment microbiota gene %GC content, and gene content of 3 *R. Alistipes* MAGs isolated from the GIT of *K. sydneyanus*. From left to right: *Bx*Car5$_{BOV}$ PUL-2, *Bx*Car17$_{BOV}$ CarPUL*,* *Bx*Car5$_{BOV}$ PUL-1. Mean %GC abundance between *Bx*Car PULs and control and marine datasets were compared using Graphpad prism. One-Way Anova with Games-Howell multiple comparisons were calculated against each PUL to identify the difference between the means. Along with previously stated controls, a native xylan PUL from each isolate was used. ns: $P > 0.05$; *: $P \leq 0.05$; **: $P \leq 0.01$; ***: $P \leq 0.001$; ****: $P \leq 0.0001$.

**Supplemental methods**

**Comparative Analysis of GC Content**

*Bx*MAG$_{BOV}$, *Bx*Car5$_{BOV}$, *Bx*Car17$_{BOV}$, control strain *B. xylanisolvens* *Bx*XB1A$_{HOM}$, and marine environment bacterium sharing the highest homology with carrageenan PUL CAZymes including *R. Alistipes* MAGs: FM_12_G117_V, FM_17_G117_V, FM_15_G124_V, *Wenyingzhuangia aestuarii* (GCF_011927765.1), *Lutibacter citreus* (GCF_003260195.1), *Labilibacter marinus* (GCF_001659685.2), *Flavobacterium* sp. 7A (GCF_025960985.1), *Saccharicrinis aurantiacus* (GCF_947489045.1), and *Saccharicrinis fermentans* (GCF_000517085.1) were annotated with prodigal (v2.6.3) to obtain GFF files. *R. Alistipes* MAGs were chosen based on their synteny with CarPULs. GC usage was calculated as a % for each gene and GraphPad Prism was used to calculate means and deviation, and Brown-Forsythe and Welch ANOVAs. Games-Howell's comparisons was done between carrageenan PULs and genomes to compare differences between GC means. Asterisks denote P value cutoff's (ns: $P > 0.05$; *: $P \leq 0.05$; **: $P \leq 0.01$; ***: $P \leq 0.001$; ****: $P \leq 0.0001$).

**Extraction of crude polysaccharide**

Ball-milled dry powder of *M. japonica* (137.6 g) was soaked in 1.2 L of hexane under magnetic stirring for 2 h in a 2 L glass beaker covered with aluminum foil to reduce evaporation. The mixture was left undisturbed overnight to allow the powder to settle. The supernatant was carefully decanted using a glass pipette, leaving approximately 1 cm of liquid above the interface to avoid disturbing the precipitate. An additional 1.2 L of hexane was added, and the extraction process was repeated. The precipitate was then resuspended in 1.2 L of 95% (v/v) ethanol, magnetically stirred for 8 h, and centrifuged at 3000 g for 30 min at room temperature. The resulting pellet was resuspended in 1.2 L of 80% (v/v) ethanol, with the same stirring and centrifugation conditions applied, and the extraction process was repeated. The resulting pellet was evaporated to dryness in 50 mL centrifuge tubes using SpeedVac (Thermo Fisher Scientific, MA, USA). The dried sample was then transferred to a glass beaker and underwent two rounds of water extraction at room temperature with constant magnetic stirring, followed by three rounds of hot water extraction at 70 °C in an incubator. Each extraction used 2 L of deionized water and lasted 8 h, with the beaker covered with aluminum foil. After each extraction, centrifugation (3,000 × *g*, 30 min, room temperature) was conducted, and the supernatant was collected while the pellet was carried forward to the next extraction. Supernatants from all the water extractions were pooled, poured into 40 L of absolute ethanol, and left at room temperature overnight, followed by centrifugation (3,000 × *g*, 30 min, room temperature). The precipitate was evaporated to dryness in 50 mL centrifuge tubes using the SpeedVac, redissolved in 2 L of deionized water by incubating at 70 °C overnight, and freeze-dried (56.7 g).

**Amylase treatment of crude polysaccharide**

Dry crude polysaccharide (14.4 g) was dissolved in 1 L of deionized water by incubating at 70 °C overnight. After that, 2 mL of thermostable α-amylase (3000 units/mL, Megazyme, Ireland) was added, and the mixture was incubated at 70 °C for 8 h [11]. The solution was then poured into 4 L of absolute ethanol, left standing at 4 °C overnight, and centrifuged (3,000 × *g*, 30 min, room temperature). The residue was evaporated to dryness using the SpeedVac, redissolved in 500 mL of deionized water by incubating at 70 °C overnight, extensively dialyzed with molecular weight cut-off (MWCO) of 6,000-8,000 Da against deionized water at 4 °C, and freeze-dried (13.8 g).

**Purification of sulfated galactan by gradient ethanol precipitation**

Amylase-treated polysaccharide (915 mg) was dissolved in 400 mL of deionized water by incubating at 70 °C overnight. The resulting solution was cooled to room temperature, left standing at 4 °C overnight, and then centrifuged (3,000 × *g*, 30 min, room temperature). The supernatant was vigorously stirred magnetically to create a water tunnel in a 2 L glass beaker. Absolute ethanol was slowly added dropwise to achieve a 15% (w/w) ethanol concentration. The mixture was kept at 4 °C for 8 h, followed by centrifugation (3,000 × *g*, 30 min, room temperature). Absolute ethanol was then added dropwise to the vigorously stirred supernatant until the ethanol concentration reached 30% (w/w), followed by standing at 4 °C and centrifugation as described above. The resulting supernatant underwent two additional cycles of ethanol precipitation, with ethanol concentrations gradually increased to 45% and 60% (w/w), respectively. The final supernatant was evaporated to dryness in 50 mL centrifuge tubes using the SpeedVac. The dry sample was redissolved in 100 mL of deionized water by incubating at 70 °C overnight, and the resulting solution was freeze-dried (0.9 g, designated as fraction F60).

**Glycosidic linkage analysis of sulfated galactan and its desulfation products**

F60 (10 mg) was dissolved in 10 mL of deionized water by magnetic stirring at 70 °C overnight, followed by 24 h of dialysis (MWCO 6,000-8,000 Da) against 4 L of 0.1 M pyridine hydrochloride, then another 24 h of dialysis against 4 L of deionized water, and freeze-dried [12]. The resulting pyridinium salt of the sulfated galactan was subjected to solvolytic desulfation by heating in 10 mL of a DMSO/methanol mixture (9:1, v/v) at 80 °C for 6 h with magnetic stirring, followed by extensive dialysis (MWCO 6,000-8,000 Da) against deionized water [8]. The sample was permethylated using 1.2 mL of methyl iodide in 2 mL of DMSO in the presence of approximately 200 mg of NaOH powder [6]. An aliquot (2 mg) of the permethylated product was subjected to reductive hydrolysis and acetylation to generate partially methylated alditol acetates (PMAAs) [13]. In another experiment, F60 was permethylated, desulfated, and converted into PMAAs in the same manner, except that the desulfation treatment lasted for 16 h instead of 6 h. In a separate experiment, F60 without desulfation treatment was changed into its triethylammonium salt form by dialysis again 0.1 M triethylamine hydrochloride, permethylated, and then converted into C-1 deuterium-labeled PMAAs by 2 M TFA hydrolysis, $NaBD_4$ reduction, and acetylation, and also into PMAAs without deuterium labeling by reductive hydrolysis and acetylation [13]. PMAA derivatives were tested on an Agilent 7890A-5977B GC-MS system (Agilent Technologies, CA, USA) equipped with a Supelco SP-2380 column (100 m × 0.25 mm × 0.2 μm; Sigma-Aldrich, MA, USA), with oven temperature programmed to start at 100 °C (hold 1 min), followed by increases of 15 °C/min to 200 °C, and then 1 °C/min to 250 °C (hold 20 min). Inlet temperature was 250°C, and the constant column helium flow rate was 1.2 mL/min. The PMAAs were identified based on their EI-MS ion fragmentation patterns [9]. Relative molar linkage compositions of the PMAAs were estimated from the total ion current (TIC) chromatogram, based on the principle that the quantity of a PMAA is proportional to the ratio of its TIC peak area to its molecular mass [14]. For each sample, two separate experiments were conducted.
